# An introduction to antibiotic-free techniques to eliminate *Staphylococcus aureus* from blood

**DOI:** 10.1101/030072

**Authors:** Gwangseong Kim, Angelo Gaitas

**Affiliations:** Kytaro, Inc., Miami, FL 33199; Florida International University (FIU), Miami, FL 33199

## Abstract

Here, we describe the implementation of three techniques for capturing and killing *Staphylococcus aureus* in blood *in vitro* inside a medical tube. The first technique involves capturing and removing pathogens using antibodies that are coated, via a simple chemical process, on the inner walls of a modified medical tube (tube capturing technique). In the second technique, a photosensitizer-antibody conjugate adheres to the pathogens while in circulation. When blood flows through the same kind of tube, the conjugate is activated by near-infrared (NIR) light to kill pathogens (photodynamic therapy technique). For the third technique, pathogens are exposed to light in the ultraviolet (UV) range while circulating through a similar tube (UV technique), which kills the pathogens.

We spiked blood with S. *aureus,* starting with about 10^7^ CFU/mL and ending at 10^8^ CFU/mL after 5 hours. While the spiked bacteria rapidly grew in nutrition-rich whole blood, each of the three techniques were able to independently remove between 61% and 84% more *S. aureus* in the experimental blood sample compared to the controls groups. When combined, these techniques demonstrated a removal rate between 87% and 92%.

## Introduction

Antimicrobial resistance is rapidly becoming a major health concern [1-3]. Resistance to antibiotics used against common pathogens, such as *S. aureus*, poses significant medical risks [4-9]. For instance, Methicillin-resistant *S. aureus* (MRSA) killed approximately 11,000 Americans in 2012 and resulted in 278,203 hospitalizations [10, 11]. The annual treatment cost associated with surgical site infections of MRSA is $12.3 billion in the USA [12]. It is evident that there is a dire need for improved treatments for patients with difficult-to-cure blood-borne infections such as MRSA.

Recently, a microfluidic device that relies on magnetic bead separation and a special bio-engineered molecule [13] was used to remove blood borne bacteria. Extracorporeal hemofiltration / hemoadsorption systems have been suggested to reduce cytokines [14] and endotoxins [15, 16] from septic blood.

Here, three antibiotic-free methodologies, which are activated while blood flows through a commercially available transparent tube, are introduced with a focus on *S. aureus* removal and killing. These techniques, shown in Fig. 1 are: (a) pathogen removal by a chemically modified medical tube coated with antibodies to capture the desired pathogens, (b) pathogen inactivation by photodynamic therapy (PDT) using photosensitizer-antibody conjugates that selectively bind to the pathogen while in circulation, allowing for the killing of these pathogens when exposed to NIR light as the blood flows through a transparent tube, and (c) UV (and more broadly near-UV) irradiation as the blood flows through the tube.

**Fig. 1.**
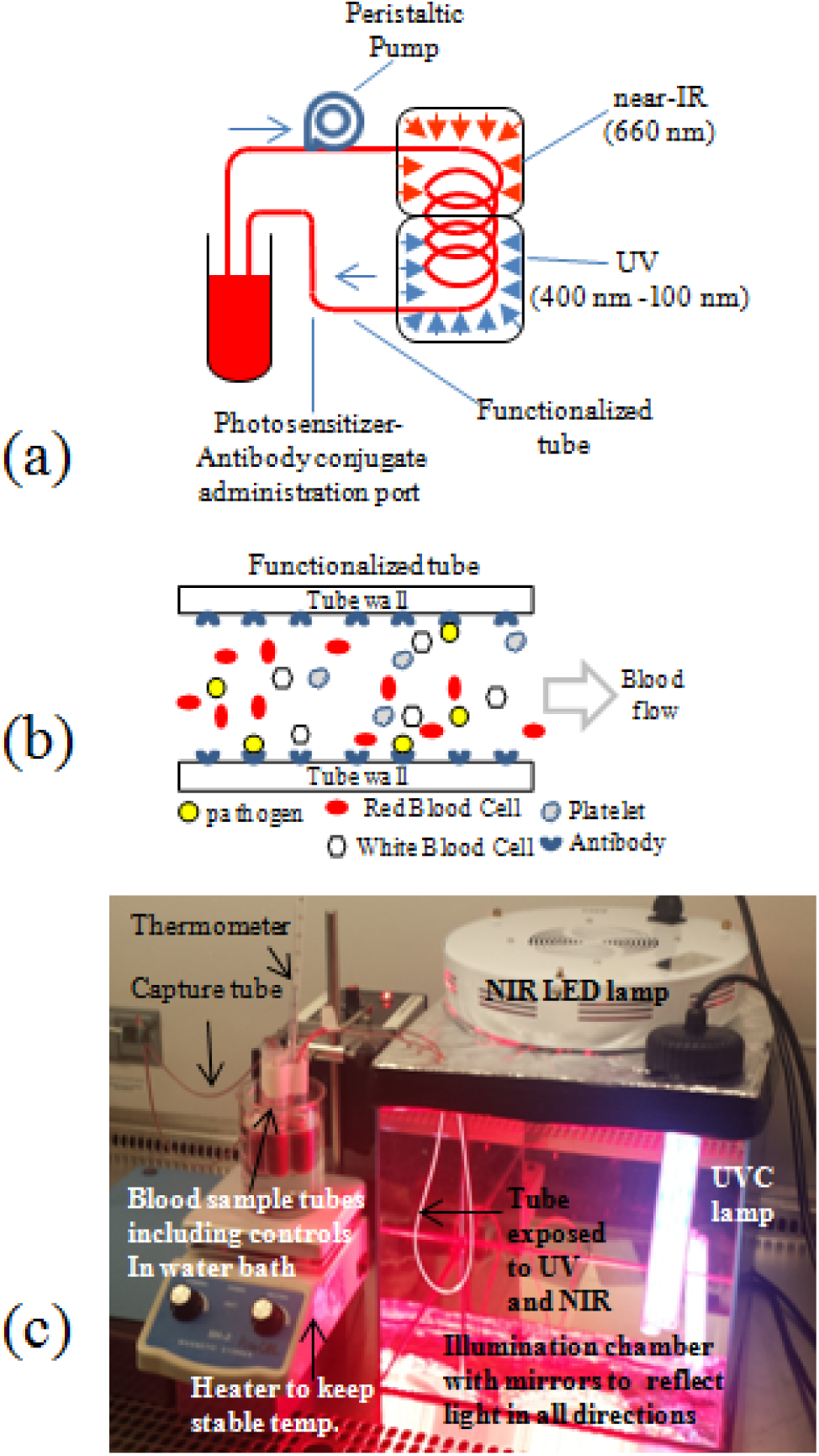
(a) Schematic of the set-up. (b) Capturing by antibody immobilization on tube. (c) Experimental setting of the system. The tubes that are exposed to UV and NIR along with the light sources are placed inside a mirror chamber. One side of the chamber was opened for this picture. The entire set-up resided inside a biosafety cabinet.

In previous efforts, we demonstrated the feasibility of the first two techniques for the removal and killing of human tumor cells [17, 18]. In this article, we describe the extension of our effort to microbial organisms in constant flow and agitation to imitate *in vivo* conditions. We also report on our investigation into the effects of combining these techniques.

## Materials and Methods

### Bacterial culture

*S. aureus* were purchased from the American Type Culture Collection (ATCC 12598). The bacteria were propagated in ATCC Medium 3 (nutrient broth or agar) at 37 °C in a shaking incubator. Bacterial concentrations were determined by both OD-600 value (optical density at 600 nm wavelength), which was measured with a UV-VIS spectrometer (Spectronic 20D+, Spectronic Instrument) in broth and its corresponding colony count from the agar plate. The initial OD-600 value in the range of 0.02-0.04 (about 1-2 × 10^7^ CFU/mL, determined by calibration). Human whole blood with an anticoagulant (sodium citrate) was purchased from vendors including Innovative Research (Novi, MI).

### Technique (a): Tube capturing

A polydimethylsiloxane (PDMS) tube (Dow Corning Silastic laboratory tubing with an internal diameter of 1.02 mm) was used for this study. The tube length was approximately 120 cm and was prepared in a manner similar to what has been described in our previous publication [18]. Specifically, the tube’s internal surface was activated by treatment with an acidic hydrogen peroxide solution (H_2_O:HCl:H_2_O_2_ in 5:1:1 volume ratio) for five minutes at room temperature [19]. The tube was rinsed with excess deionized (DI) water five times and dried in air. The tube was then filled with aminopropyltrimethoxysilane (APTMS) for 10 minutes. The tube was rinsed with excess DI water at least five times and dried in air.

*S. aureus* polyclonal antibody (PA1-7246, Life Technologies) was treated for 1.5 hour with Traut’s reagent (2-iminothiolane HCl, 2-IT) to generate an available sulfhydryl group (-SH) (antibody:2-IT=1:10 in mole ratio) in PBS (pH 7.4). Then, unbound 2-IT was removed from the antibodies using a protein concentrator (MWCO 30 kDa, Corning Spin-X protein concentrator) at 5000 RCF for 30 minutes. The concentrated *S. aureus* pAb was re-suspended in PBS, and the volume was adjusted to 1.2 mL. During the antibody-2-IT reaction, the amine functionalized tube was filled with a hetero-bifunctional crosslinker, sulfo-SMCC (sulfosuccinimidyl 4-[N-maleimidomethyl]cyclohexane-1-carboxylate) in 2 mg/mL concentration in PBS (pH 7.4). After the 2-IT treated pAb was spun down, the sulfo-SMCC solution was removed, and the tube was rinsed in PBS and re-filled with 1.2 mL re-suspended *S. aureus* pAb solution. The reaction was run on a shaker for two hours at room temperature and continued overnight at 4 °C. The next day, after the unbound antibody solution was collected, the tube was gently rinsed with PBS and then refilled with 2 mg/mL L-cysteine for another two hours. The tube was rinsed and filled with PBS and stored at 4 °C until use.

0. 5 mL of *S. aureus* (10 × concentrated) were added to 4.5 mL of whole blood, resulting in 5 mL of 1× bacterial solution in blood (estimated bacterial input was 1-2 × 10^7^ CFU/mL) in sterile 15 mL culture tube. The initial bacterial concentration was estimated by monitoring the OD-600 value of sample diluted in the same way in broth instead of blood. Blood-bacteria mixtures were stirred with a mini magnetic stirrer to prevent blood from settling down by gravity over time. *S. aurues pAb* immobilized tubes were connected to the blood-bacteria mixture with both ends submerged into the solution.

### Technique (b): PDT

2 mg of a NIR photosensitizer, Chlorin E6 (Ce6, Frontier Scientific) were mixed with 6.5 mg of 1-Ethyl-3-[3-dimethylaminopropyl]carbodiimide hydrochloride (EDC, crosslinker) (Sigma-Aldrich) and 7.6 mg of sulfo-NHS (stabilizer for EDC) (Pierce) in 1 mL 10% Dimethyl sulfoxide - PBS buffer (DMSO:PBS=10:90), (Ce6:EDC:sulfo-NHS=1:10:10 in mole ratio). The reaction was run at room temperature with agitation for 2 hours. Then, 1 mL of 200 μg/mL *S. aureus* pAb in 10% DMSO-PBS mixture was mixed with 1 mL of Ce6 mixture. The conjugation reaction was run at room temperature with agitation for 3 hours. The reaction mixture was spin-filtered with a protein concentrator to remove the unbound Ce6 and other chemicals from the desired Ce6-antibody conjugates at 5000 RCF for 15 min, and the procedure was repeated 4 times with refilling excess 10% DMSO-PBS solution. The final product was re-suspended in PBS, adjusting the final volume of 1 mL. The produced Ce6-conjugated *S. aureus* pAb was stored at 4 °C. All conjugates were consumed within 1-2 weeks from their preparation.

### Technique (c): UV (or near UV) exposure of extracorporeal tube

Its germicidal effects have made UV light a valuable tool for killing bacteria and viruses. UV irradiation has been used in surgical wound disinfection with high success rates for eliminating bacteria [20, 21]. Here, *S. aureus* spiked blood samples were circulated through an unmodified PDMS tube, which was inserted into an illumination chamber. UV light (ODYSSEA UVC-18W, centered at 254 nm) was used to illuminate the tube inside the illumination chamber.

### Combination of multiple techniques

Three individual techniques were combined in different configurations of two and three. The combinations used were: PDT-tube capturing (using 1 capturing tube and 1 unmodified tube), UV-tube capturing (using 1 capturing tube and 1 unmodified tube), and PDT-UV-tube capturing (using 1 capturing tube and 1 unmodified tube for both PDT and UV). In all combinations, each tube was directly connected to one blood sample in a parallel connection. Tubes for PDT, UV, or both were inserted into illumination chambers with NIR LED lamps and UV lamps installed on top.

### Experimental setup

The experimental setup included three major components: a temperature-controlled bath, a peristaltic pump, and an illumination chamber. *S. aureus* spiked blood samples were placed in 15 mL sterile culture tubes and in a water bath heated to 37^o^ C on a small heating stirrer plate. In PDT, the Ce6-pAb conjugate (200 µL) was added to the blood. The blood sample was agitated with a mini magnetic stirrer (7 mm × 2 mm) inside the solution. A 120 cm tube for techniques (a), (b), and (c) was connected to the blood sample in the water bath. The tube was installed through a peristaltic pump (P-3, Pharmacia) to maintain a constant flow of blood. Part of the tube was inserted into a cube-shape illumination chamber (12 × 12 × 12 inch) with mirror walls to reflect the light. The temperature inside the illumination chamber was monitored and tested for several hours to ensure that heat generated by the 660 nm LED lamb and the UV lamp did not reach temperatures that could cause thermal damage to cells. The chamber temperature was equilibrated in the range of 29 - 31^o^ C. The tubes were connected to the blood samples in a water bath to complete the circulation system. The entire set up (shown in Fig 1 (c)) was installed inside a biosafety cabinet to prevent contamination. The *S. aureus* containing blood mixture was circulated through the tube with peristaltic pumping at 0.5 mL/min for 5 hours. In the PDT case, in order to allow sufficient time for the antibody conjugates to bind with *S. aureus*, the blood mixture was circulated through the tube without illumination for the first 2 hours. Illumination by NIR light was then performed for 3 hours.

### Processing samples and colony counting

*S. aureus* was pipetted from a culture tube and diluted until the value fell between 1 10^7^ and 2 10^7^ CFU/mL (initial OD-600 0.02-0.04). We included a control each time the experiment was performed to account for the variation in bacteria concentration and growth patterns between stocks (deviation from stock to stock).

During the procedure, 50 μL blood samples were extracted at the following time intervals: 0, 1, 3, and 5 hours. The sample was diluted in 450 μL of nutrient broth (10 × dilution) and diluted 3 more times in same manner (100 ×, 1000 ×, 10000 × dilution). 10 μL of each of the diluted samples were streaked on a 5% sheep blood agar plate (Fisher Scientific) that was divided into quadrants. The bacteria colonies were allowed to grow overnight. The colonies were then counted using the particle analysis function in ImageJ (National Institutes of Health). Each experiment and control was repeated 5 times.

## Results

Technique (a), tube capturing, yielded bacterial growth 11.4 ± 4.3 × 10^7^ CFU/mL (mean ± standard error of mean (SEM), n=5), while the control growth (from same stock) was 60.5 ± 16.3 × 10^7^ CFU/mL at 5 hour as shown in Fig. 2 (a). Using only PDT, there was a reduction of 10.9 ± 3.0 × 10^7^ CFU/mL, whereas the control group produced a reduction of 40.9 ± 10.1 × 10^7^ CFU/mL (Fig. 2 (b)). UV tube exposure was the least effective of the techniques when applied alone with 32.8 ± 4.2 × 10^7^ CFU/mL vs. 83.8±2.5 × 10^7^ CFU/mL for the control (Fig. 2 (c)). When PDT and tube capturing were combined, there was a reduction in bacteria of 9.2 ± 3.5 × 10^7^ CFU/mL vs. 69.4 ± 8.31 × 10^7^ CFU/mL for the control (Fig. 2 (d)). When combining UV and tube capturing the CFUs were 9.0 ± 2.7 × 10^7^ CFU/mL vs. 80.4 ± 4.3 × 10^7^ CFU/mL for the control (Fig. 2 (e)). Finally, when all three techniques were combined, the experimental group’s reduction in bacteria was 4.2 ±1.0 × 10^7^ CFU/mL, compared to 59.7 ± 2.5 × 10^7^ CFU/mL for the control group (Fig. 2 (f)).

**Fig. 2.**
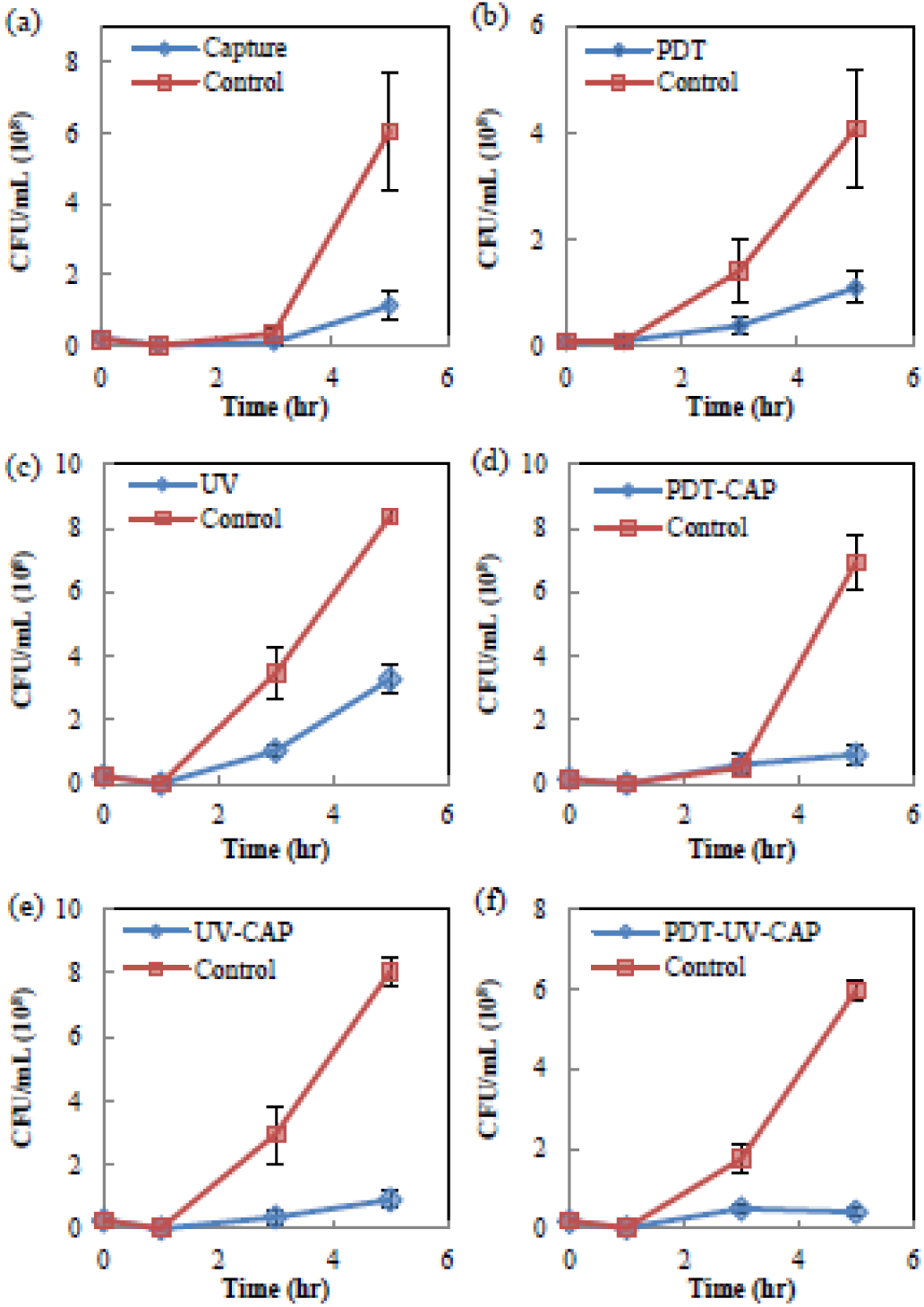
Graph of bacterial growth using the techniques described vs. control after 5 hours (n=5) (CFU/mL; error bars, SEM.), (a) Capturing (t-test P<0.02), (b) PDT (t-test P<0.03), (c) UV (t-test P < 0.0001), (d) PDT-Capturing combination (t-test P < 0.0002), (e) UV-Capturing combination (t-test P < 0.0001), and (f) PDT-UV-Capturing combination (t-test P < 0.0001).

## Discussion

Given that the initial bacteria count was not exactly the same for the control and the experiment groups, the values in Fig. 3 were normalized before comparison with controls. Using tube capturing, there were 83.4% fewer bacteria, while using PDT there were 71.4% fewer bacteria. UV light yielded a 61.6% bacteria reduction. Among the three individual techniques, capturing appeared to be the most effective for removing bacteria. Combining these techniques yielded improved suppression of *S. aureus* growth. When PDT and tube capturing were combined, there was a reduction in bacteria CFU by 87.1%. When UV and tube capturing were combined, the percent reduction of the bacteria CFU was 89%. Finally, when all three techniques were combined, the experimental group’s reduction in bacteria was 92%.

**Fig. 3.**
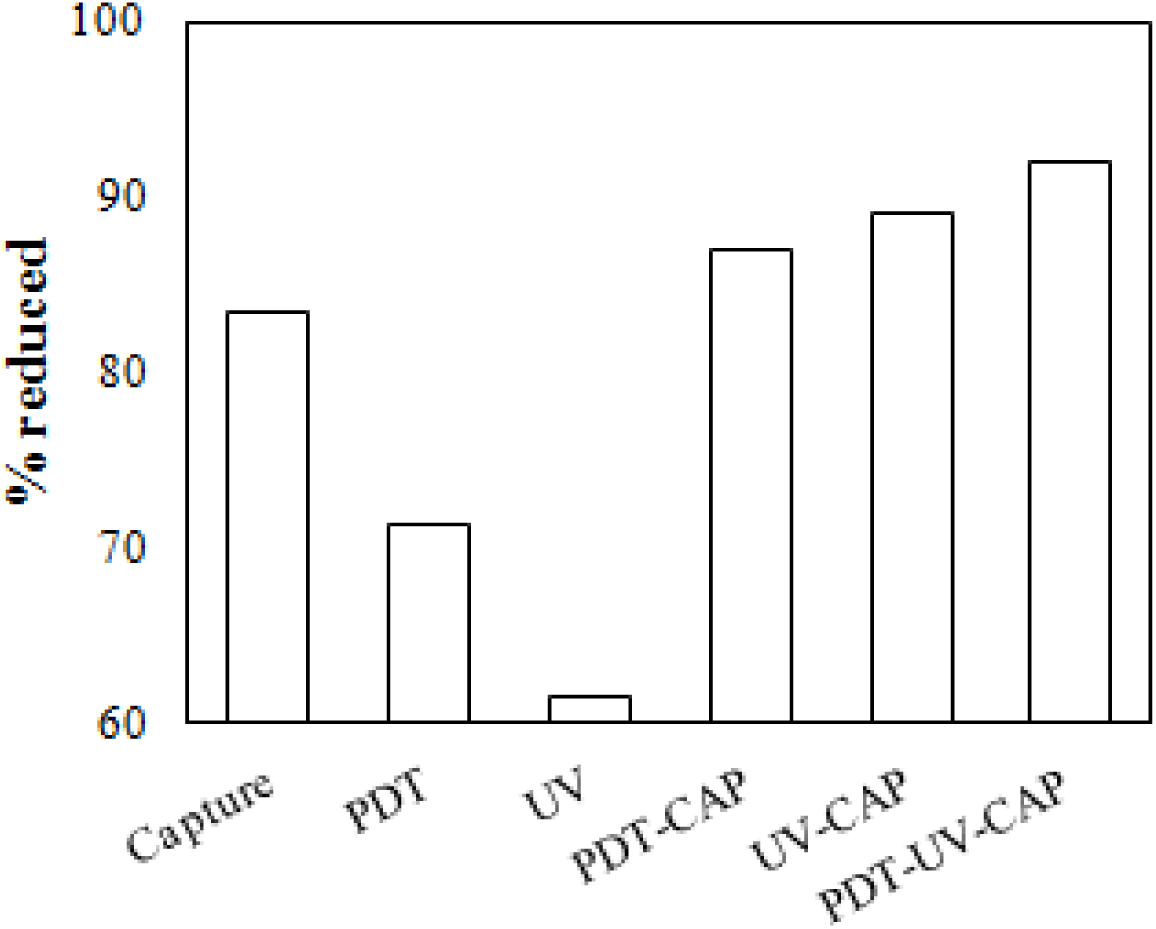
Summary of results of normalize values in % compared to control.

When combined, these techniques reduce bacterial growth at the given bacteria concentrations *in vitro* with higher efficiency than individual techniques. These conclusions are dependent on the initial bacteria concentration, type of bacteria, and organism. The utilization of a thin plastic transparent tube provides the foundation of these three techniques. Tube capturing does not kill bacteria, so using this technique alone will likely lead to bacteria repopulation. Thus, the two additional methods were employed to further inactivate the pathogens.

The utilization of a thin tube was essential for PDT to function in blood. PDT is based on the activation of photosensitizers by light. The presence of hemoglobin in blood (a strong light absorber) blocks the majority of light to achieve effective PDT. PDT has been used in applications where deep tissue penetration by light is not required (e.g. skin cancer [22, 23], lung [24, 25], head and neck cancer [26, 27], and some dental conditions [28, 29]). Our set-up allowed PDT to function successfully in blood by utilizing: (a) a thin transparent tube that allows the entire blood sample volume flowing through the tube to be exposed to NIR light, and (b) a NIR photosensitizer with an excitation wavelength of 660 nmm where the light absorption by hemoglobin is minimal.

PDT’s efficacy is based on oxidative damage by locally induced reactive oxygen species. PDT efficacy depends on oxidative stress, which is non-specifically effective within narrow vicinities. Thus, PDT can kill antibiotic-resistant microorganisms, such as MRSA [30, 31]. The photosensitizer must be selectively delivered by conjugating with an adhesion molecule to target organisms to prevent collateral damage to other blood components.

The presence of excess and unbound photosensitizers (Ce6-pAb in this case) in the blood may cause non-specific damage to blood cells. Preliminary studies by other groups indicate that PDT in blood “is safe *in vivo*” [32, 33]. Potential toxicity will be investigated in the future given that toxicity investigation is more relevant when this technology is advanced enough for animal studies in which the bone marrow and the body’s filtering organs create a more realistic scenario. Current clinical practices of PDT require a waiting period to minimize undesired collateral damage [34]. This period occurs between the application of the photosensitizer and the light illumination to allow for the photosensitizer to accumulate in target cells/tissues and for the excess photosensitizer to be cleared by the body through its natural filtering mechanisms. Such clearance cannot be emulated in *in vitro* conditions. The dosage of the photosensitizer and appropriate waiting time will have to be carefully determined by further studies to maximize the PDT’s efficacy on targeted pathogens and at the same time to minimize adverse effects from unbound photosensitizers in circulation.

Another concern regarding toxicity is collateral damage to distant cells by reactive oxygen species’ (ROS) diffusion/convection from targeted pathogens, which is highly unlikely. The predominant ROS in photosensitization is the molecular singlet oxygen (^1^O_2_), which is extremely short lived and has very limited free diffusion distance (reportedly, 0.01-0.02 μm) in biological media [35, 36]. Also, prior research has demonstrated that PDT’s efficacy is strictly confined to targeted cells in the mixture of different cells [37]. Future toxicity studies will analyze targeted versus non-targeted cell death with such studies as complete blood count (CBC) differential test, applying separate fluorescent tags, or radiolabels, in addition to a cell viability assay and analyzing cell death with a cell sorting technology.

Utilizing thin tubes enables the use of UV in blood. The germicidal effect of UV light is well established and has been widely used in sterilization, including sterilization of wounds. Health risks by UV, including carcinogenesis, have restricted the use of UV light in humans. Here, UV irradiation was performed only on tubes residing in a mirrored chamber. Hemoglobin in blood significantly blocks light in the UV range, thus the resulting penetration depth of UV would be even shorter than that of PDT [38, 39]. Therefore, it seems that bacterial killing by UV likely occurs very close to the tube’s inner surface, which may explain the lower efficiency of this technique compared to capturing and PDT. The appeal of using germicidal light lays in the non-specific damage mechanism that allows for effectively eliminating most microorganisms. Potential toxicity by UV irradiation on normal blood components will be investigated in future work. However, it is worth noting that recent findings have shown that wavelengths in the low 400 nm are also capable of disinfecting pathogens, including *S. aureus* and MRSA [40-43] and that short wavelength UV rays (UVC, 207 nm) can selectively kill bacteria with negligible damage to mammalian cells [20]. Further investigation will be conducted exploring various wavelengths to effectively inactivate pathogens without health risk.

There are a number of ways that these techniques can be further improved. For instance, introducing an additional antibody or binding molecule may further enhance efficiency. In this work, capturing and PDT both used the same antibody to bind to *S. aureus.* Introducing an additional binding molecule could reduce a potential competition for binding cites. The technology presented here is currently limited in its capacity to handle high volumes of blood efficiently. Various engineering optimizations are underway to address conditions for larger animals and humans.

The capturing tube and the photosensitizer-antibody conjugates can be easily prepared with a specific antibody. Adhesion molecules that target a large group of pathogens will be included in future experiments, eliminating the need to first identify the pathogen. These general-purpose molecules can be used to coat the tube and create the conjugate. For instance, antibodies or molecules adhering to galactose-alpha-1,3-galactose (alpha gal), a carbohydrate found in the cell membrane of most organisms, but not in human cells, can be used as a target.

In addition to removing *S. aureus*, these techniques can be applied to remove Gram-positive and Gram-negative bacteria, fungi, viruses, parasites and other types of microorganisms using appropriate adhesion molecules. These techniques can be used to reduce infectious particle load to minimal levels or at levels where conventional medication and the body’s own immune system can fight the infection. Thus, they may be particularly useful for individuals experiencing immunosuppression or young children for whom antibiotics and antifungal medication can be highly toxic [44, 45]. Extending these concepts further, future work will include tubes coated with pathogen-killing agents. It will also interesting to use the antibody capturing technique as an enrichment device. By circulating blood through a series of capturing tubes, each with a specific antibody (or other targeting molecules), microorganisms can be concentrated in the tubes without requiring further isolation steps. This approach could reduce the time required to identify a pathogen.

## Conclusion

Three antibiotic-free techniques that can be adapted to reduce or eliminate bacterial infections from blood have been presented. In this manuscript, we used *S. aureus* and a combination of techniques to reduce bacteria load by about 90% *in vitro* in spiked blood. These techniques could be adaptable for use against other microorganisms and possibly against antimicrobial resistant strains. Several procedures should be implemented to improve these techniques, such as testing additional microorganisms, answering engineering challenges like throughput, and studying the toxicity of PDT and germicidal light.

## Acknowledgments

We would like to thank Drs. Aileen Marty, Yogesh Gianchandani, Roma Gianchandani, Shekhar Bhansali, Horatiu Vinerean, and Paula Flicker for their feedback, comments, and support.

## References

1. Barfod TS, Wibroe EA, Brauner JV, Knudsen JD. Changes in antimicrobial susceptibility patterns of Klebsiella pneumoniae, Escherichia coli and Staphylococcus aureus over the past decade. Dan Med J. 2015;62(10). Epub 2015/10/07. PubMed PMID: 26441391.

2. Prestinaci F, Pezzotti P, Pantosti A. Antimicrobial resistance: a global multifaceted phenomenon. Pathogens and global health. 2015:2047773215Y. 0000000030.

3. Bush K, Courvalin P, Dantas G, Davies J, Eisenstein B, Huovinen P, et al. Tackling antibiotic resistance. Nature Reviews Microbiology. 2011;9(12):894–6.

4. Rhee Y, Aroutcheva A, Hota B, Weinstein RA, Popovich KJ, editors. Evolving Epidemiology of Staphylococcus aureus Bacteremia. IDWeek 2014; 2014: Idsa.

5. Deresinski S. Methicillin-resistant Staphylococcus aureus: an evolutionary, epidemiologic, and therapeutic odyssey. Clinical infectious diseases. 2005;40(4):562–73.

6. Gomes A, Westh H, De Lencastre H. Origins and evolution of methicillin-resistant Staphylococcus aureus clonal lineages. Antimicrobial agents and chemotherapy. 2006;50(10):3237–44.

7. Deurenberg R, Vink C, Kalenic S, Friedrich A, Bruggeman C, Stobberingh E. The molecular evolution of methicillin - resistant Staphylococcus aureus. Clinical Microbiology and Infection. 2007;13(3):222–35.

8. Harris SR, Feil EJ, Holden MT, Quail MA, Nickerson EK, Chantratita N, et al. Evolution of MRSA during hospital transmission and intercontinental spread. Science. 2010;327(5964):469–74.

9. Grema H, Geidam Y, Gadzama G, Ameh J, Suleiman A. Methicillin resistant Staphyloccus aureus (MRSA): a review. Adv Anim Vet Sci. 2015;3(2):79–98.

10. Klein E, Smith DL, Laxminarayan R. Hospitalizations and deaths caused by methicillin-resistant Staphylococcus aureus, United States, 1999-2005. Emerging infectious diseases. 2007;13(12):1840.

11. National strategy for combating antibiotic resistant bacteria 2014. Available from: https://www.whitehouse.gov/sites/default/files/docs/carb_national_strategy.pdf.

12. Anonymous. Press Release: Pfizer Begins Phase 2b Study Of Its Investigational Multi-antigen Staphylococcus aureus Vaccine In Adults Undergoing Elective Spinal Fusion Surgery. 2015.

13. Kang JH, Super M, Yung CW, Cooper RM, Domansky K, Graveline AR, et al. An extracorporeal blood-cleansing device for sepsis therapy. Nature medicine. 2014.

14. Zhang J, Peng Z, Maberry D, Volpe J, Kimmel J, Federspiel W, et al. Effects of Hemoadsorption with a Novel Adsorbent on Sepsis: In vivo and in vitro Study. Blood purification. 2015;39(1–3):239–45.

15. Shoji H. Extracorporeal Endotoxin Removal For The Treatment of Sepsis:Endotoxin Adsorption Cartridge (Toraymyxin). Therapeutic Apheresis and Dialysis. 2003;7(1):108–14. doi: 10.1046/j.1526-0968.2003.00005.x.

16. Peng Z-Y, Zhang J, Rimmelé T, Zhou F, Chuasuwan A, Kaynar AM, et al. Development of venovenous extracorporeal blood purification circuits in rodents for sepsis. Journal of Surgical Research. 2013;185(2):790–6.

17. Kim G, Gaitas A. Extracorporeal photo-immunotherapy for circulating tumor cells. PloS one. 2014;10(5):e0127219-e.

18. Gaitas A, Kim G. Chemically Modified Plastic Tube for High Volume Removal and Collection of Circulating Tumor Cells. PLoS One. 2015;10(7):e0133194. doi: 10.1371/journal.pone.0133194.

19. Sui G, Wang J, Lee C-C, Lu W, Lee SP, Leyton JV, et al. Solution-phase surface modification in intact poly (dimethylsiloxane) microfluidic channels. Analytical chemistry. 2006;78(15):5543–51.

20. Buonanno M, Randers-Pehrson G, Bigelow AW, Trivedi S, Lowy FD, Spotnitz HM, et al. 207-nm UV light-a promising tool for safe low-cost reduction of surgical site infections. I: in vitro studies. PloS one. 2013;8(10):e76968.

21. Ritter MA, Olberding EM, Malinzak RA. Ultraviolet lighting during orthopaedic surgery and the rate of infection. The Journal of Bone & Joint Surgery. 2007;89(9):1935–40.

22. Baldea I, Filip A. Photodynamic therapy in melanoma–an update. J Physiol Pharmacol. 2012;63(2):109–18.

23. Kawczyk-Krupka A, Bugaj AM, Latos W, Zaremba K, Sieroń A. Photodynamic therapy in treatment of cutaneous and choroidal melanoma. Photodiagnosis and photodynamic therapy. 2013;10(4):503–9.

24. Simone CB, II JSF, Glatstein E, Stevenson JP, Sterman DH, Hahn SM, et al. Photodynamic therapy for the treatment of non-small cell lung cancer. Journal of thoracic disease. 2012;4(1):63.

25. Allison R, Moghissi K, Downie G, Dixon K. Photodynamic therapy (PDT) for lung cancer. Photodiagnosis and photodynamic therapy. 2011;8(3):231–9.

26. Schuller DE, McCaughan JS, Rock RP. Photodynamic therapy in head and neck cancer. Archives of Otolaryngology. 1985; 111(6):351–5.

27. Vesper BJ, Colvard MD. Photodynamic Therapy (PDT): An Evolving Therapeutic Technique in Head and Neck Cancer Treatment. Head & Neck Cancer: Current Perspectives, Advances, and Challenges: Springer; 2013. p. 649–76.

28. Gursoy H, Ozcakir-Tomruk C, Tanalp J, Yilmaz S. Photodynamic therapy in dentistry: a literature review. Clinical oral investigations. 2013; 17(4): 1113–25.

29. Silva LAB, Novaes AB, de Oliveira RR, Nelson-Filho P, Santamaria M, Silva RAB. Antimicrobial photodynamic therapy for the treatment of teeth with apical periodontitis: a histopathological evaluation. Journal of endodontics. 2012;38(3):360–6.

30. Dai T, Tegos GP, Zhiyentayev T, Mylonakis E, Hamblin MR. Photodynamic therapy for methicillin - resistant Staphylococcus aureus infection in a mouse skin abrasion model. Lasers in surgery and medicine. 2010;42(1):38–44.

31. Fu X-j, Fang Y, Yao M. Antimicrobial photodynamic therapy for methicillin-resistant Staphylococcus aureus infection. BioMed research international. 2013;2013.

32. Yin H, Zhang G, Chen H, Wang W, Kong D, Li Y. Preliminary safety evaluation of photodynamic therapy for blood purification: an animal study. Artificial organs. 2014;38(6):510–5.

33. Yin H, Ye X, Li Y, Niu Q, Wang C, Ma W. Evaluation of the effects of systemic photodynamic therapy in a rat model of acute myeloid leukemia. Journal of Photochemistry and Photobiology B: Biology. 2015.

34. https://www.roswellpark.org/patients/treatment-services/innovative-treatments/photodynamic-therapy.

35. Kadish KM, Smith KM, Guilard R. Handbook of Porphyrin Science: With Applications to Chemistry, Physics. Materials Science, Engineering, Biology and Medicine. 2010;1.

36. Kihara S, Hartzler DA, Savikhin S. Oxygen concentration inside a functioning photosynthetic cell. Biophysical journal. 2014;106(9):1882–9.

37. Avula UMR, Kim G, Lee Y-EK, Morady F, Kopelman R, Kalifa J. Cell-specific nanoplatform-enabled photodynamic therapy for cardiac cells. Heart Rhythm. 2012;9(9):1504–9.

38. Takatani S, Graham MD. Theoretical analysis of diffuse reflectance from a two-layer tissue model. Biomedical Engineering, IEEE Transactions on. 1979;(12):656–64.

39. Schmitt J. Optical Measurement of blood oxygenation by implantable telemetry. Technical Report, 1986.

40. Maclean M, MacGregor SJ, Anderson JG, Woolsey G. High-intensity narrow-spectrum light inactivation and wavelength sensitivity of Staphylococcus aureus. FEMS microbiology letters. 2008;285(2):227–32.

41. Dai T, Gupta A, Huang Y-Y, Yin R, Murray CK, Vrahas MS, et al. Blue light rescues mice from potentially fatal Pseudomonas aeruginosa burn infection: efficacy, safety, and mechanism of action. Antimicrobial agents and chemotherapy. 2013;57(3): 1238–45. doi: doi: 10.1128/AAC.01652-12. Epub 2012 Dec 21.

42. Lipovsky A, Nitzan Y, Gedanken A, Lubart R. Visible light-induced killing of bacteria as a function of wavelength: Implication for wound healing. Lasers in surgery and medicine. 2010;42(6):467–72. doi: 10.1002/lsm.20948.

43. Lubart R, Lipovski A, Nitzan Y, Friedmann H. A possible mechanism for the bactericidal effect of visible light. Laser therapy. 2011;20(1): 17.

44. Hassan M, Yasmeen BN, Begum N. Fungal sepsis and Indications of antifungal prophylaxis and treatment in neonatal intensive care units: A review. Northern International Medical College Journal. 2015;6(1):6–8.

45. Murray MJ. Critical care medicine: perioperative management: Lippincott Williams & Wilkins; 2002.

